# Structural Genomic Variation And Its Potential Role In Deer Speciation

**DOI:** 10.1101/2025.04.23.650217

**Authors:** Faezeh Azimi Chetabi, Aaron Shafer

**Affiliations:** 2140 East Bank Drive, Environmental and Life Sciences Graduate Program, Trent University, Peterborough, Ontario, Canada K9J 7B8

**Keywords:** Comparative genomics, short-read, long-read, adaptation, genomic divergence, *Odocoileus*

## Abstract

Speciation is a key driver of biodiversity and understanding its genomic underpinnings can be important for predicting and managing biodiversity. Structural variants (SVs) are large-scale (>50 bp) changes in the genome and have been implicated in adaptive divergence and reproductive isolation. We investigated the role of SVs in the speciation and divergence of two deer species (*Odocoileus* spp.) across their North American range. Using multiple long-read and a short-read datasets, our bioinformatics workflow revealed SVs and genomic features that were unique to each species. The majority of species-specific SVs were deletions and insertions suggesting that these variants may show higher likelihoods of fixation within populations. Further, while most SVs were intergenic, some genes were impacted, with 3 species-specific SVs showing signs of selection inferred from dN/dS. We also observed a reduced number of regulatory motifs found in fixed SVs compared to the rest of the genome. The SV-affected genes were often associated with reproduction and sensory adaptation, with such functions being relevant to fertility and deer biology and therefore providing insights into potential mechanisms leading to reproductive divergence.

## 1. Introduction

Speciation is a fundamental process driven most notably by ecological divergence and geographic isolation (Sobel, 2010), with the ultimate mode of divergence leaving tell-tale signals on the genome (Shang et al., 2023). Structural variants (SVs) have long been considered important in driving divergence, with early cytogenetic work pointing to chromosomal differences as barriers to reproduction (Rieseberg, 2001). In mammals for example, two distinct clades of Brocket deer (*Mazama* spp.) show karyotypic differences that contribute to reproductive isolation (Abril et al., 2010). Similarly, in donkey-horse hybrids (*Equus* spp.), failed chromosomal pairing during spermatogenic meiosis leads to sterility (Chandley et al, 1974; Chandley et al 1975). More recently, comprehensive reviews have highlighted that SVs, ranging from inversions to translocations, can influence speciation across diverse taxa (Wellenreuther et al., 2019; Berdan et al., 2023). The advent of high-throughput sequencing and novel bioinformatics tools has made it possible to assay SVs at base-pair resolution, opening a new line of inquiry into their role in adaptation and speciation across virtually all taxa.

While single nucleotide polymorphisms have historically dominated evolutionary genomics, SVs by definition affect larger stretches of DNA and can encompass entire genes and regulatory regions and may have stronger fitness consequences (Wellenreuther et al. 2019). Structural variants are generally considered features >50 bp and fall into several main types: deletions, duplications, inversions, insertions and translocations. Different sequencing and analytical techniques can be employed to detect SVs. Short-read (SR) data with paired read and split read analyses offer high-resolution breakpoint detection, though their effectiveness may depend on factors like insert size and coverage depth (Escaramís et al., 2015). Read depth analysis, often used with either SR or long-read (LR), provides information into SV presence by examining read density, but is less precise with breakpoint resolution (Escaramís et al., 2015). For LR data, de novo assembly and genome alignments are frequently employed, and offers the advantage of identifying novel SVs without relying on reference genomes, though it is more resource-intensive (Escaramís et al., 2015). Long-read sequencing is generally preferable than SR sequencing for detecting complex and large SVs, as SR methods struggle with novel insertions and complex rearrangements (Mahmoud et al., 2019). More broadly, LR sequencing is proving transformative, revealing the prevalence and complexity of SVs across genomes and reshaping how we study divergence (Berdan et al., 2023).

Structural variants such as inversions and translocations can promote speciation through mechanisms like reproductive isolation (Homolka et al., 2007). Recombination suppression by SVs can also couple adaptive alleles and incompatibilities, reinforcing divergence even in the face of gene flow (Berdan et al., 2023; Wellenreuther et al., 2019). Autosomal translocations have been shown to cause male sterility in laboratory mice due to disrupted synapsis and meiotic silencing (Homolka et al., 2007), while Yang et al., (2024) showed convergent evolutionary signals of SVs associated with domestication, adaptation, and various traits. Further, Gompert et al. (2025) demonstrated that complex SVs, including inverted translocations, can independently arise in separate populations and repeatedly drive adaptive divergence by suppressing recombination and clustering functionally important genes, revealing a predictable mechanism of local adaptation and evolutionary change. Structural variants can also drive phenotypic diversity by influencing gene expression (Zhang et al, 2024), with enhancers often connecting genetic changes to species-specific phenotypes (Kaplow et al., 2023) and diseases (Fudenberg et al 2019; also referred to as enhancer-hijacking). Shi. et. al (2023) presented some evidence for enrichment of promoters and enhancers in specific SV categories. Collectively, these findings highlight SVs and enhancer regions as potential drivers of evolutionary divergence, shaping unique phenotypes and enabling evolution of traits across species.

The *Cervidae* (Deer) family offers an interesting case study for exploring SVs and genetic history. This family encompasses over 60 species with its origins dating back nearly 20 million years (Gonzalez and Duarte, 2020). The red brocket deer (*Mazama americana*) exhibits at least six distinct karyotypes across South America, suggesting the presence of multiple cryptic species (Cursino et al., 2014). Studies on hybrids between these karyotype lineages revealed post-zygotic reproductive isolation, with hybrids showing sterility or subfertility due to chromosomal imbalances impacting ovarian structure, oocyte development, and embryonic genome activation (Cursino et al., 2014). Two closely related species, white-tailed deer (WTD; *Odocoileus virginianus*) and mule deer (MD; *O. hemionus*), have a relatively recent speciation history, appearing to have split in allopatry just over 1 million years ago (Ma) but also undergoing contemporary hybridization (Kessler et al., 2023). Kessler et al., (2023) suggested the existence of genetic incompatibilities given the lack of nuclear introgression and historical gene flow. Collectively, this raises the possibility that SVs might have played a role in deer speciation and genomic divergence.

In this study, we integrated multiple LR and SR sequencing datasets from MD and WTD across their geographic ranges to identify and characterize species-specific SVs. This comparative framework, which combines diverse sequencing approaches and detection tools (Figure 1), enables a comprehensive assessment of the evolutionary impact of SVs. We hypothesized that fixed SVs are important to deer speciation between the two deer species through two, non-mutually exclusive mechanisms: by driving adaptive divergence through changes in genic and regulatory regions, and by generating genomic incompatibilities that reduce hybrid fitness and reinforce reproductive isolation. By identifying consistent, species-specific SVs and examining their overlap with functional genomic regions and regulatory motifs, this study aims to better understand how SVs shapes adaptation, divergence, and the formation of species boundaries in *Odocoileus*.

**Figure 1.**
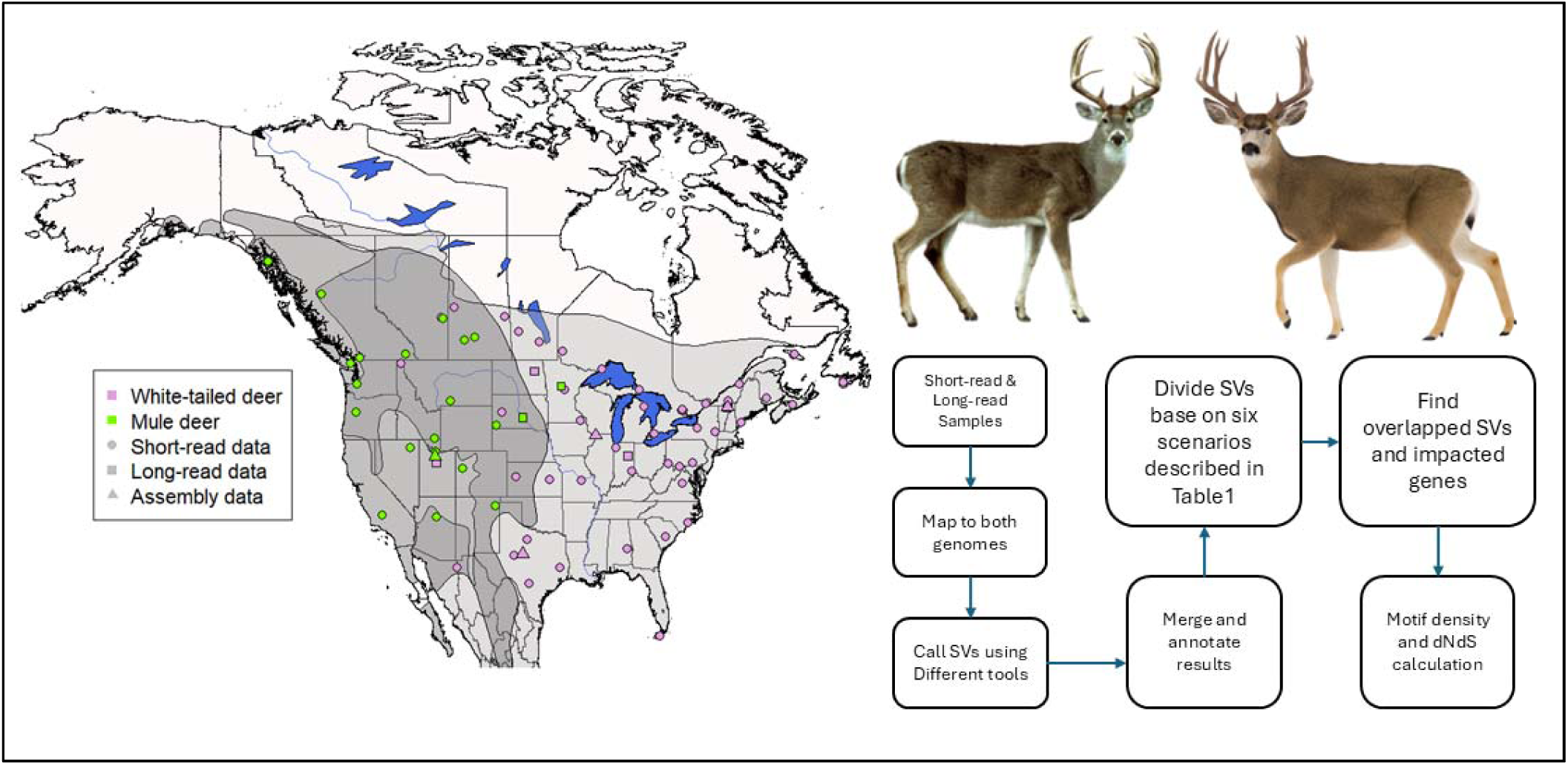
General workflow of this study depicted on the right. The map shows the geographical range of the two species under investigation. The light grey area represents the range of white-tailed deer, while the grey area indicates the range of mule deer. The dark grey region highlights the overlap between the two species’ habitats. Each point indicates the specific locations where samples were collected.

## 2. Materials and Methods

### 2.1. Data generation and retrieval

We collated publicly available Illumina whole-genome short-read datasets from 79 deer (57 WTD and 22 MD; BioProject PRJNA830519); DNA was extracted from tissue samples (Qiagen DNeasy Blood and Tissue Kit) and sequenced at the Centre for Applied Genomics (Toronto, Canada) on an Illumina HiSeqX platform (see Kessler et al., 2023 for more information). We also retrieved two LR datasets from NCBI (SRR6668252 for WTD and SRR15720448 for MD).

We generated new LR data from five deer semen samples (3 WTD and 2 MD); samples were first thawed on ice and cells were pelleted in an isotonic sperm wash solution to remove debris and reduce somatic cell contamination. DNA was extracted using the NEB Monarch® HMW DNA Extraction Kit for Cells & Blood (#T3050) UHMW protocol, with some modifications to the cell lysis step. Specifically, cells were digested at 56°C for 1 hour at 300 rpm with 100 μL Nuclei Prep Buffer, 100 μL Nuclei Lysis Buffer, 10 μL Proteinase K (NEB #P8107, 20 mg/ml), and 10 μL 1M DTT (GoldBio #DTT in dH2O, final concentration ∼50 mM). An additional 20-minute digestion was then performed with 5 μL RNase A (NEB #T3018, 20 mg/ml). DNA quality control, CCS library preparation, and sequencing on the PacBio Revio sequencing instrument were then conducted. Collectively, there are four LR samples for WTD and three LR samples for MD. General characteristics and quality of the long-read datasets used in our analyses were assessed using SeqKit (Shen et al., 2016). For each LR sample, we summarized the total number of reads, and the minimum, average, and maximum read lengths.

We acquired two reference genome assemblies from the National Center for Biotechnology Information (NCBI) database: one of WTD (JAJQKH000000000) and the other of MD (JAJLRB000000000). To assess the completeness of these assemblies, we evaluated them using BUSCO (Simão et al, 2015), with the score of 98.3% for WTD and 98.0% for MD. We used the GENESPACE v1.1.4 R package (Lovell et al., 2022), that required OrthoFinder (Emms et al, 2019), DIAMOND (Buchfink et al, 2021) and MCScanX (Wang et al, 2012), to identify syntenic blocks and visualize macrosynteny relationships between WTD (*Odocoileus virginianus*), MD (*Odocoileus hemionus*), Caribou (*Rangifer tarandus*, CATKSN000000000), and domestic cattle (*Bos taurus*, NKLS00000000) (omitting *Cervus* for alignment compatibility reasons). To visualize chromosomal relationships, we generated riparian (macrosynteny) plots using the plot_riparian() function in GENESPACE. Color palettes and themes were customized using the R language. The WTD genome was used as the reference in the visualization.

### 2.2. SV detection pipeline

We employed two distinct analytical approaches to detect SVs (Figure 1). We used both LR datasets and SR data in conjunction with highest quality genome per species to detect species specific SVs (Figure S1-S2). Because reference genome assemblies can produce artefacts (Howe et al, 2021), and we recognized that there are multiple sequencing data and reference genome combinations, we identified the scenarios that generated species-specific SVs (Table 1). Specifically, the SVs detected in scenarios 4 and 5 are relatively unambiguous as species-specific SV. Likewise, scenarios 3 and 6 (both con-specific mapping) are likely only to arise via an assembly artefact or an SV singleton and are therefore not species-specific. Scenarios 1 and 2 are SVs detected with both reference assemblies, but not the sister species resequencing data; here we consider the SV to be real, and the SV is absent or misassembled in the reference. All SV indels are defined relative to the reference genome, meaning each reported indel represents reference-relative classifications rather than absolute evolutionary events.

**Table 1.**
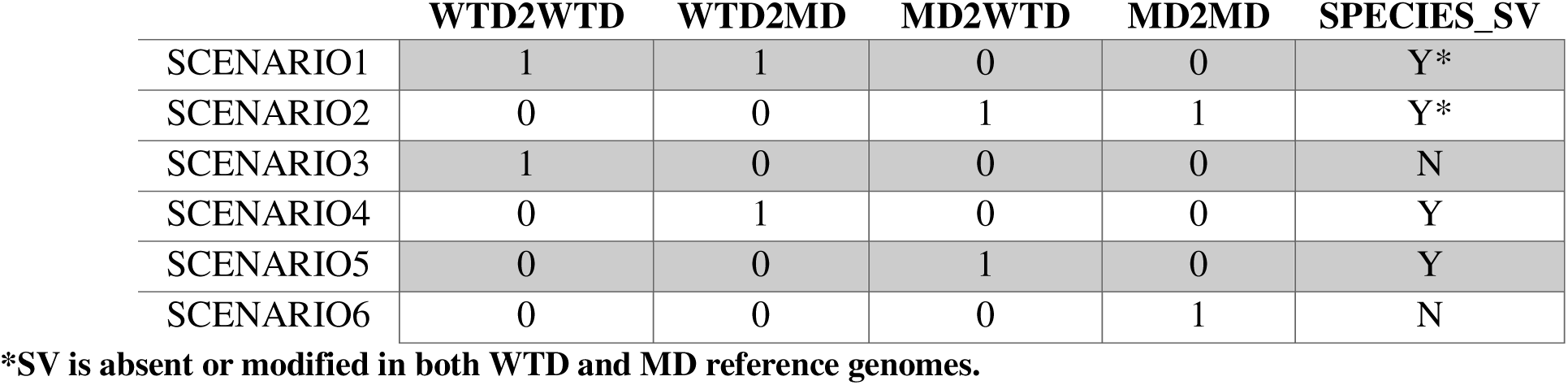
Summary of the mapping scenarios used to identify species-specific structural variants (SVs) in both short-read and long-read datasets. WTD and MD refer to white-tailed deer and mule deer, respectively. The first four columns indicate the mapping strategy for each dataset—that is, which species’ samples were aligned to which reference genome, with 1 meaning all samples have the SV. For example, “WTD2WTD” represents white-tailed deer samples mapped to the white-tailed deer reference genome. Last column indicates if the SVs derived from that strategy were considered species-specific (Y) or no (N).

|  | WTD2WTD | WTD2MD | MD2WTD | MD2MD | SPECIES_SV |
| --- | --- | --- | --- | --- | --- |
| SCENARIO1 | 1 | 1 | 0 | 0 | Y* |
| SCENARIO2 | 0 | 0 | 1 | 1 | Y* |
| SCENARIO3 | 1 | 0 | 0 | 0 | N |
| SCENARIO4 | 0 | 1 | 0 | 0 | Y |
| SCENARIO5 | 0 | 0 | 1 | 0 | Y |
| SCENARIO6 | 0 | 0 | 0 | 1 | N |

\*SV is absent or modified in both WTD and MD reference genomes.

For the SR pipeline, we removed adaptors from the raw fastq files using Trimmomatic (Bolger et al., 2014) with a MINLEN threshold of 36. We employed Bowtie2 (Langmead et al., 2012) to align all samples to both the reference genomes of WTD and MD. We used SAMtools (Danecek et al., 2021) to convert, sort, and index the SAM files. To ensure data integrity, duplicate reads were eliminated using Picard (Broad Institute, 2018) with REMOVE_SEQUENCING_DUPLICATES set to TRUE. We employed three different SV calling tools for SR data: Delly (Rausch et al., 2012), Manta (Chen et al., 2016), and Lumpy (Layer, 2014), all with default settings. Further refinement and comparative analysis of VCFs were performed using BCFtools (Danecek et al., 2021), retaining only structural variants with SUPP > 5, representing the number of sequencing reads supporting a given SV. We merged the SV call sets from Delly, Manta, and Lumpy using Jasmine (Kirsche et al, 2023), configured to confirm SVs found in at least two tools. Using BCFtools (Danecek et al., 2021), we filtered the structural variants to extract specific information for comparison, such as the number of SVs, their sizes in base pairs (bp) and zygosity status.

Our LR analysis pipeline followed a workflow similar to that described by Mérot et al. (2023). First, we mapped the LR datasets to the WTD and MD reference genomes using Winnowmap (Jain et al, 2020). The aligned SAM files were then converted to BAM format, sorted, and indexed with SAMtools (Danecek et al., 2021). Structural variants were identified using a combination of Sniffles (Sedlazeck et al., 2018), SVIM (Heller et al, 2019), and NanoVar (Tham et al, 2020). Finally, we employed Jasmine (Kirsche et al, 2023) to merge the SV calls into a single VCF file. We filtered the merged VCF files based on read support greater than 5.

Both LR and SR pipelines produced four mapping categories: MD2MD, MD2WTD, WTD2WTD, and WTD2MD (Figure S1-S2). For each individual, SVs were classified according to whether they were detected in: (1) both WTD2WTD and WTD2MD; (2) both MD2WTD and MD2MD; (3) only WTD2WTD; (4) only WTD2MD; (5) only MD2WTD; or (6) only MD2MD (Table 1). This classification framework enabled us to identify species-specific SVs (See Table 1, Figure S1-S2). The SVs from scenarios 3 and 6 were not included in further analysis,

To allow cross-referencing of features between species, we used MUMmer (Delcher et al., 2002) to align the WTD and MD reference genomes. Using the *show-coords* utility, we extracted the corresponding aligned intervals and identified regions shared between the two genomes and accurately lift over scaffold coordinates. To further refine our dataset and identify SVs with the potential relevance to speciation, we retained only those SVs present in 100% of the individuals within each scenario. In practice, this meant requiring SVs be detected in all 22 MD SR samples, or in all 57 WTD SR samples, or in all 4 WTD LR samples, or in all 3 MD LR samples, depending on the scenario. This filtering generated species-specific VCF datasets for both WTD and MD across the LR and SR data (Figure S1-S2). Finally, we quantified the overlap of these species-specific SVs between LR and SR analyses within each scenario.

### 2.3. Gene Impact, Selection Analysis and Enhancer Motifs

We annotated the MD and WTD genome for enhancer motifs using Homer (Heinz et al., 2010). The findMotifsGenome, annotatePeaks, and scanMotifGenomeWide tools were employed and filtered based on the average motif scores. We extracted the number of motifs detected and compared the number in intergenic regions, to those within 50 kb of any SV, and within 50 kb of species-specific SVs (up- and downstream); here, we normalized motif density, by dividing the number of motifs by the length of each region to produce a value per kilobase pair (kbp) and compared groups using a ANOVA. The resulting impact of SVs from both LR and SR analysis was annotated using SnpEff (Cingolani et al., 2012). To enable this, we created custom SnpEff databases using the genome FASTA files and their corresponding annotation GFF files. As with cross-referencing genomes, we transferred annotations between assemblies using Mummer (Delcher et al., 2002) and custom Python scripts.

Lastly, we calculated the ratio of nonsynonymous to synonymous substitution rates (dN/dS) for specific genes overlapping and the fixed (high confidence) SV categories; we prepared coding sequences (CDS) and corresponding protein sequences for the same species used for synteny analysis (omitting *Alces* for alignment compatibility reasons). Single copy orthologous genes were identified using OrthoFinder (Emms et al, 2019) with DIAMOND (Buchfink et al, 2021) for sequence similarity searches, MAFFT (Katoh et al., 2005) for multiple sequence alignment, and FastTree (Price etal, 2009) for species tree inference. For each orthogroup, corresponding CDS sequences were extracted using seqtk (Li, 2012) and aligned at the codon level using PAL2NAL (Suyama et al, 2006). Gene sequences were used as input for BUSTED (Murrell et al, 2025) method implemented in HyPHy (Kosakovsky Pond et al., 2005) to estimate dN/dS ratios. The predefined species tree topology (Cow, (Elk, (Caribou, ((MuleDeer, WhiteTailedDeer))))); was used in all analyses. Evidence of selection on individual MD or WTD lineages using the foreground analysis was assessed using likelihood ratio tests with significance determined at p < 0.05 after Bonferroni-Holm correction.

## 3. Results

### 3.1. Genome SV exploration

The Cervidae genomes appear to be largely syntenic at a macro-level (Figure 2). Average sequencing coverage for the SR datasets was approximately 6X; the corresponding LR datasets showed variation in coverage, but all were > 7X (Table 2). Structural variant distribution across all individuals in the SR and LR merged SV datasets (i.e. 2/3 methods detected the SV) are shown in Figure 3 for mapping to the conspecific genome (i.e. WTD2WTD and MD2MD). Here, the median SV size in bp presented WTD then MD for deletion was 206 and 239, duplication was 1817 and 2027, insertion was 84 and 89 bps, inversion was 1294 and 2501 bps and translocation was 610 and 850; average individual SV homozygosity was >50% for indels (Table S1). In the LR merged SV dataset, the median SV size in bp presented WTD then MD for deletion was 136 and 133, duplication was 9821 and 5531, insertion was 109 and 132 bps, inversion was 570 and 1178 bps and translocation was 201 and 746; average individual SV homozygosity was 30%-70% for indels (Table S1).

**Figure 2.**
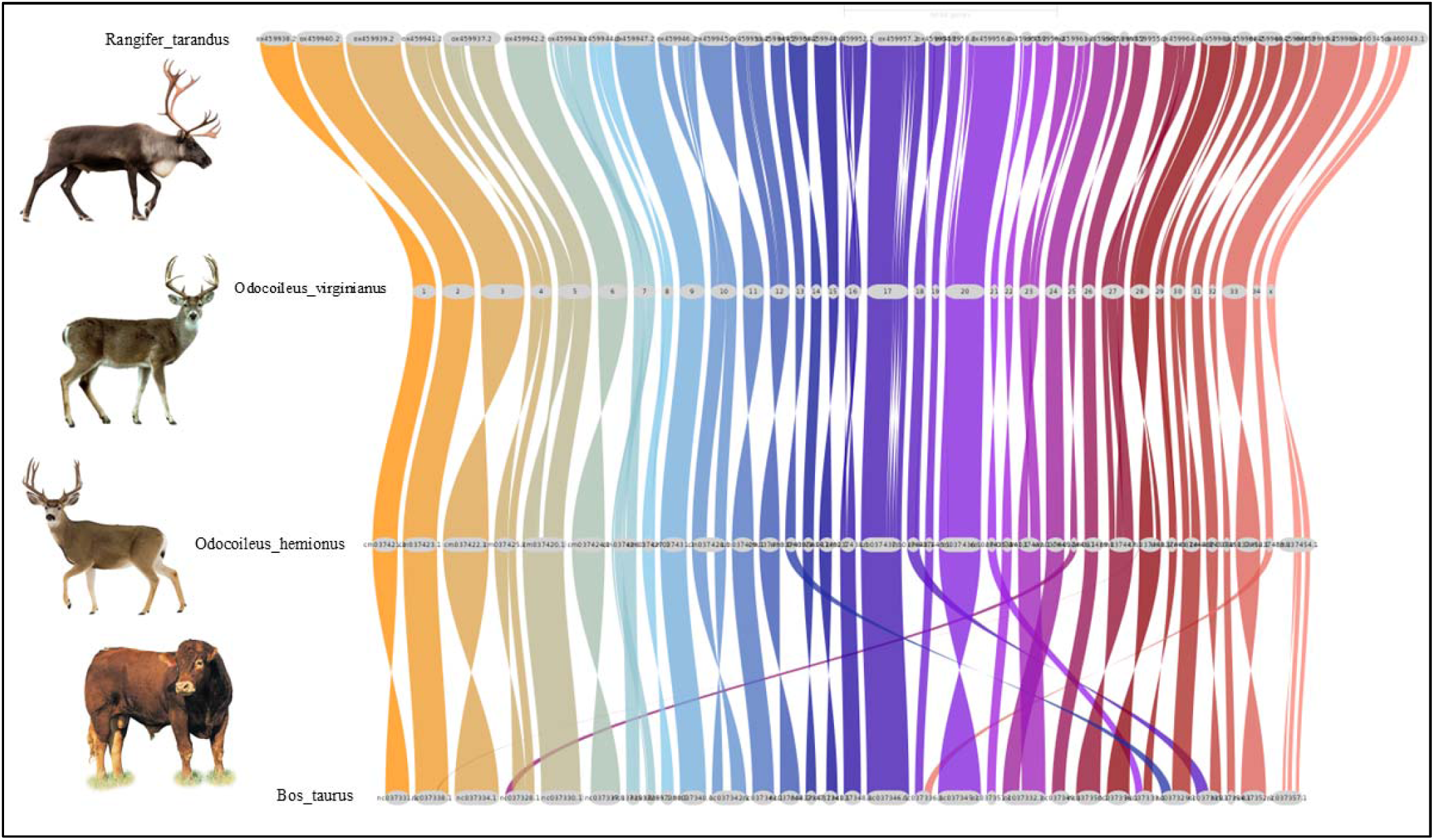
Macrosynteny comparison among white-tailed deer (*Odocoileus virginianus*), mule deer (*Odocoileus hemionus*), caribou (*Rangifer tarandus*), and domestic cattle (*Bos taurus*). Chromosome segments are connected by colored ribbons, showing conservation of chromosome structure and highlighting the close genomic relationship between white tailed and mule deer.

**Table 2.**
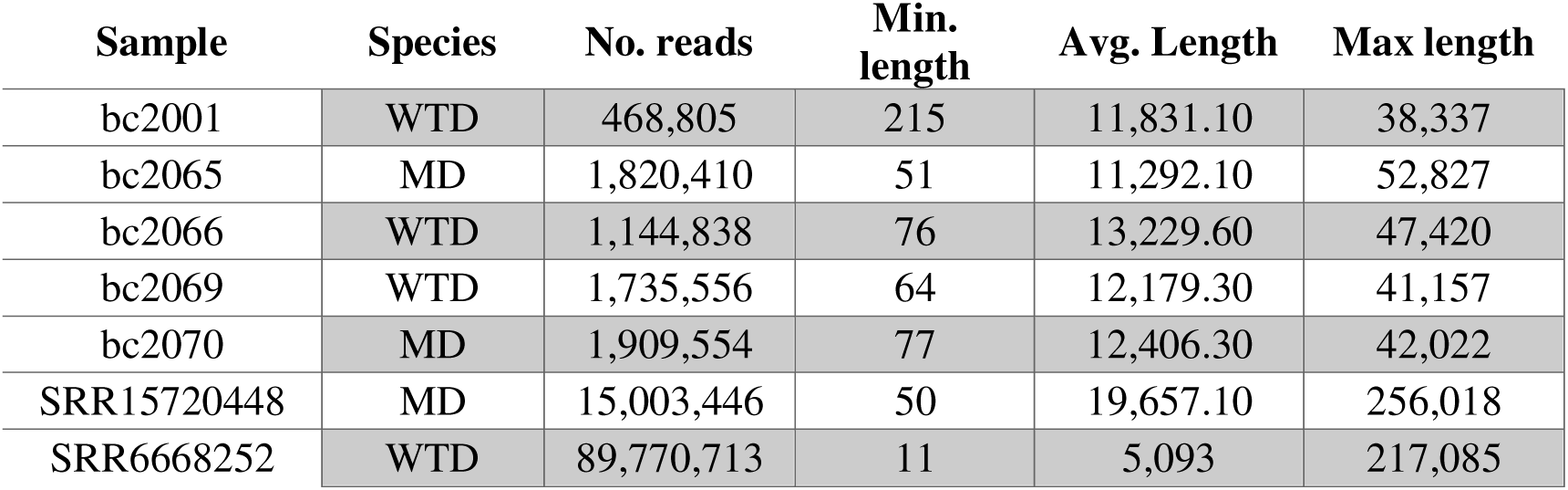
Raw sequencing statistics for all long-read samples from white-tailed deer (WTD) and mule deer (MD). For each dataset, the total number of reads (No. reads), minimum read length (Min. length), average read length (Avg. Length), and maximum read length (Max length) are reported.

| Sample | Species | No. reads | Min.<br>length | Avg. Length | Max length |
| --- | --- | --- | --- | --- | --- |
| bc2001 | WTD | 468,805 | 215 | 11,831.10 | 38,337 |
| bc2065 | MD | 1,820,410 | 51 | 11,292.10 | 52,827 |
| bc2066 | WTD | 1,144,838 | 76 | 13,229.60 | 47,420 |
| bc2069 | WTD | 1,735,556 | 64 | 12,179.30 | 41,157 |
| bc2070 | MD | 1,909,554 | 77 | 12,406.30 | 42,022 |
| SRR15720448 | MD | 15,003,446 | 50 | 19,657.10 | 256,018 |
| SRR6668252 | WTD | 89,770,713 | 11 | 5,093 | 217,085 |

**Figure 3.**
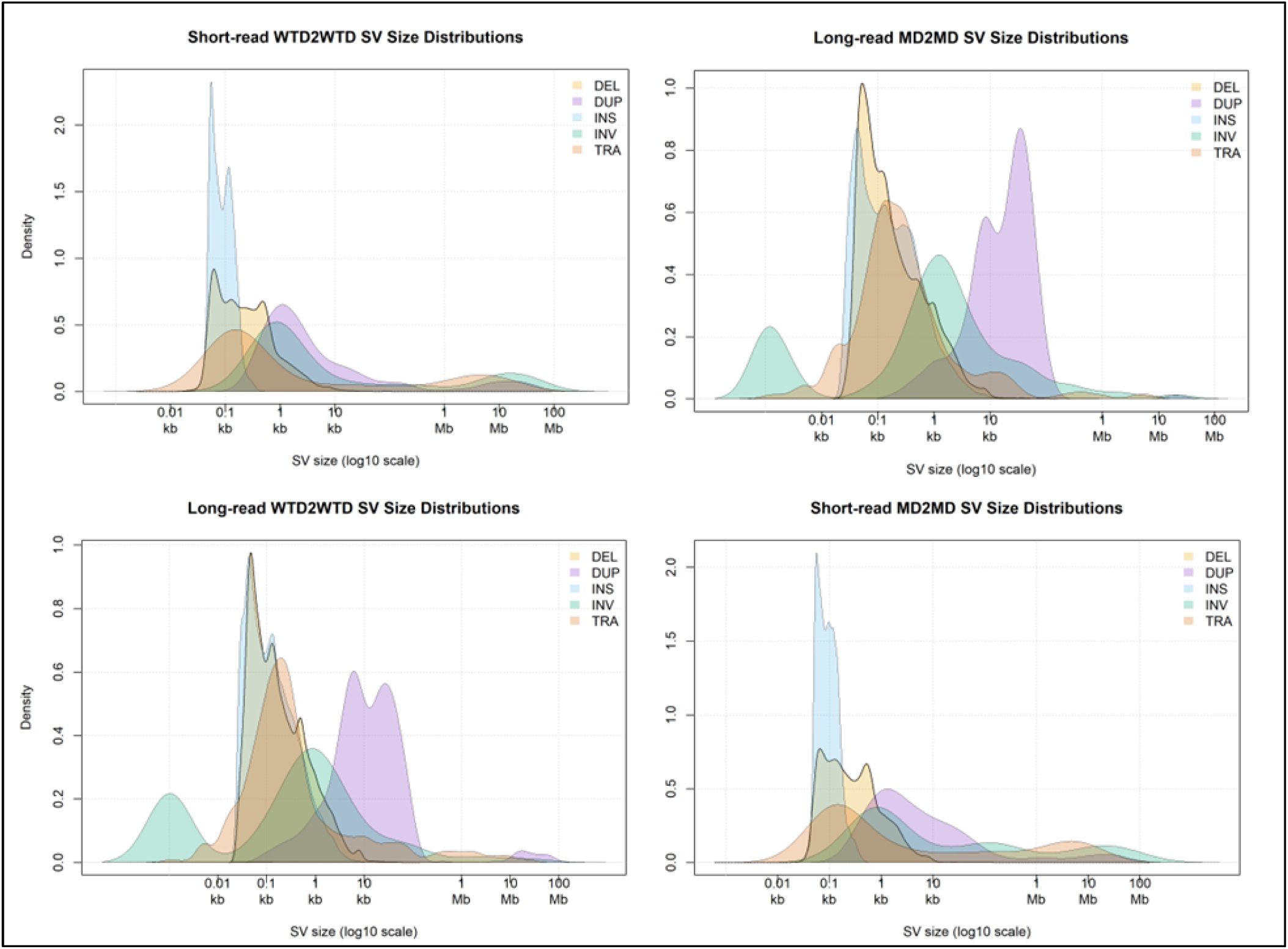
Comparison of the number of structural variants (SVs) detected across SV types in the entire short read dataset. Each panel represents a different mapping strategy, showing the counts of deletions (DEL), duplications (DUP), insertions (INS), inversions (INV), and translocations (TRA) on a log scale.

### 3.2. Species specific SVs, Genome and Enhancer Impact

We next focused on species-specific SVs that were present in 100% of samples within each species in Scenarios 1,2,4,5 (Table 1). In total we identified 56 unique SVs in 100% of SR samples (Table 3). In contrast, we identified 123,384 unique SVs in 100% of LR samples (Table 4). The majority of predicted gene-impacts were intronic (Table 3 and 4). We assessed the overlap between SR- and LR-derived SVs by considering only those variants located within exonic, intronic, or gene-level regions (Table 5); intergenic SVs were excluded from this comparison. Of note 3 of genes had significant dN/dS values in WTD and MD; we identified notable genes in fixed SVs as it pertains to deer biology and potential reproductive isolation (Table S2). For both MD and WTD, enhancer motifs were elevated in intergenic regions, with those adjacent to fixed SVs having the lowest density (Table S3; ANOVA F = 45.25, p < 0.01).

**Table 3.** Species-specific structural variants (SVs) detected in short-read dataset across the scenarios defined in Table 1, using the criterion that each SV must be present in 100% of samples. For each scenario, the table reports: (i) the total number of SVs; (ii) counts of each SV class (DEL, DUP, INS, INV, TRA); (iii) the number of genes overlapping those SVs; and (iv) the distribution of gene impacts, categorized as exonic, intronic or gene-level. SV types are abbreviated as follows: DEL = deletion, DUP = duplication, INS = insertion, INV = inversion, TRA = translocation.

| SR - 100% of samples |  |  |  |  |
| --- | --- | --- | --- | --- |
|  | # SVs | SV Types | # Genes | Gene Impact |
| Scenario1 | 11 | DEL:0 DUP:0 INS:3 INV:0 TRA:8 | 11 | 0\3\8 |
| Scenario2 | 10 | DEL:1 DUP:0 INS:0 INV:1 TRA:8 | 9 | 0\0\9 |
| Scenario4 | 4 | DEL:2 DUP:0 INS:1 INV:0 TRA:1 | 3 | 0\0\3 |
| Scenario5 | 31 | DEL:13 DUP:6 INS:0 INV:2 TRA:10 | 17 | 4\1\12 |

**Table 4.** Species-specific structural variants (SVs) detected in the long-read dataset across the scenarios defined in Table 1, using the criterion that each SV must be present in 100% of samples. For each scenario, the table reports: (i) the total number of SVs; (ii) counts of each SV class (DEL, DUP, INS, INV, TRA); (iii) the number of genes overlapping those SVs; and (iv) the distribution of gene impacts, categorized as exonic, intronic, or gene-level. SV types are abbreviated as follows: DEL = deletion, DUP = duplication, INS = insertion, INV = inversion, TRA = translocation.

**LR - 100% of samples**
|  | # SVs | SV Types | # Genes | Gene Impact |
| --- | --- | --- | --- | --- |
| *Scenario1* | 19280 | DEL:8341 DUP:0 INS:10801 INV:138 TRA:0 | 4371 | 241\4127\3 |
| *Scenario2* | 14552 | DEL:7911 DUP:99 INS:4156 INV:217 TRA:2169 | 6220 | 426\3570\2224 |
| *Scenario4* | 23074 | DEL:14646 DUP:0 INS:8381 INV:47 TRA:0 | 2316 | 79\2050\187 |
| *Scenario5* | 66478 | DEL:37002 DUP:0 INS:29295 INV:181 TRA:0 | 12134 | 1307\10716\111 |

**Table 5.**
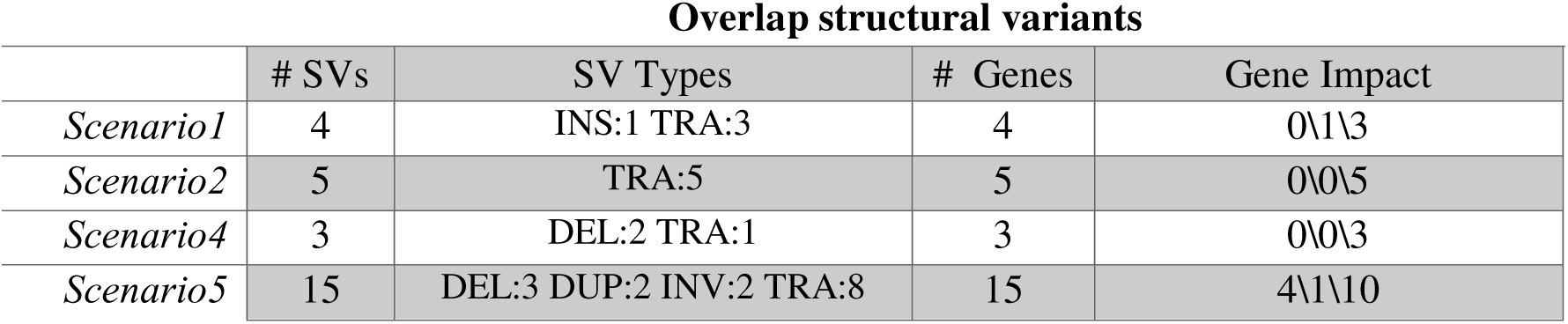
Species-specific structural variants (SVs) detected in both long and short read datasets across the scenarios defined in Table 1. For each scenario, the table reports: (i) the total number of SVs; (ii) counts of each SV class (DEL, DUP, INS, INV, TRA); (iii) the number of genes overlapping those SVs; and (iv) the distribution of predicted gene impacts, categorized as exonic, intronic, or gene-level. SV types are abbreviated as follows: DEL = deletion, DUP = duplication, INS = insertion, INV = inversion, TRA = translocation.

## 4. Discussion

The genomes of recent and hybridizing species provide a window into the mechanisms that drive biodiversity, adaptation, and reproductive isolation. We compared both SR and LR data on two closely related and hybridizing deer species, the white-tailed deer and mule deer, to identify SVs fixed between species. By integrating data from multiple LR and SR sequencing, we identified SVs within and between these species, their predicted impact on genes, and general relationship to regulatory motifs. Genome analyses has detected negligible historical introgression between these species, despite contemporary hybridization (Kessler et al., 2023), suggesting that much of the observed genomic divergence reflects drift and isolation rather than ongoing exchange. This lack of gene flow complicates the identification of barrier loci, since patterns of divergence cannot be readily distinguished from neutral processes, but there is very little evidence for any meaningful nuclear gene-flow (Kessler et al. 2023). While the presence of species-specific SVs reflects major evolutionary events, we acknowledge that fixed differences alone cannot demonstrate a causal role in reproductive isolation, as divergence also accumulates neutrally in the absence of gene flow. Nonetheless, the fixed SVs that encompass genes and regulatory regions provides testable hypotheses and clues to species divergence and reproductive isolation.

### General characteristics of SVs in *Odocoileus*

Our analysis revealed a predominance of insertions and deletions among SVs in deer. This finding is consistent with observations reported in other studies (Shi et al., 2023; Zhang et al., 2023), notably, Shi et al. (2023) identified deletions as the most frequent SV type in human populations. This pattern could be attributed to various mechanisms, including replication errors or unequal homologous recombination (Wold et al., 2023). Berdan et al., (2023) summarized how SVs, particularly when heterozygous, can suppress recombination, but homozygous deletions and insertions, by their nature, do not inherently disrupt the linear arrangement of genetic material, minimizing their impact on recombination. Duplications and translocations, particularly in their heterozygous form, can result in complex genomic rearrangements that disrupt gene order and heighten the risk of recombination errors. Zhang et al. (2023) discussed how gene duplications can drive speciation by disrupting gene function or fostering novel interactions, with empirical evidence showing duplications clearly underlying incompatibilities (Zuellig et al 2018; Bikard et al 2009). While the species-specific duplications were among the least common SV we detected (Table 3 and 4), these are worth further functional exploration, especially those containing genic regions.

Most of the SV impacts we observed were intergenic, and similar trends were reported by Yang et al. (2024) and David et al. (2024). These studies, like ours, demonstrated that a significant portion of SVs localized to intergenic regions in sheep, goats, and wild bird populations. This dominance of intergenic impacts can be attributed to the vast amount of non-coding DNA within the genome; however, these regions also contain important regulatory elements, which comprise a significant portion of the mammalian genome (Villar et al, 2015). David et al. (2024) observed that a high percentage of deletions and inversions in intergenic regions were classified as modifiers, suggesting they could have phenotypic effects. Our findings provide a similar perspective where 80.11% of deletions and 72.99% of inversions occurred in intergenic regions and 18.44% of intergenic deletions and 15.99% of intergenic inversions were annotated as modifiers, reiterating the potential consequential impacts SVs can have outside genic regions. The reduced density of enhancers in SVs, particularly those in fixed SVs (Table S3), highlight the influence of intergenic space in the genome when it comes to promoters and enhancers (ENCODE Project Consortium, 2012). Shi et al. (2023) found a genome-wide positive correlation between enhancer and promotors and SV density. The reduction in motifs in the fixed SVs is an interesting observation in deer; while enhancer and silencer action are often cell dependent (Panigrahi et al, 2021), this pattern is consistent with a positional or cis-ruption effect (see Kleinjan and Coutinho, 2009), meaning the regulatory architecture is impacted, with the molecular and phenotypic impact in deer warranting further investigation.

### Species-species SVs, adaptation and potential incompatibilities

The identification of fixed SVs highlights potential connections to adaptive divergence. Recent studies in Eucalyptus species (Ferguson et al., 2024) showed the role of shared SVs in adaptation and speciation with significant number of SVs being enriched for genes involved in adaptive traits. Likewise, work on stick-insects showed clustering functionally important genes (Gompert et al. 2025). In deer, patterns were nuanced but did align with predictions and previous work, with deletions being the most common fixed SV. This can be functionally important as Derks et al. (2018) showed in pigs that a large homozygous deletion that exhibited the highest penetrance. In slight contrast, Redin et al. (2017) found that inversions and translocations van have stronger association congenital abnormalities. Accordingly, the higher rate fixed deletions between species might point to a role in adaptive divergence and speciation.

Several presumed gene impacts were identified that align with known aspects of deer biology and presumed speciation (Kessler et al., 2024). Notably, there were multiple genes with known links to male gonad development and sperm production (Table S2), which is important because hybrid deer show evidence of reduced sperm function (Derr et al., 1991). Similarly, both Bracewell et al. and Dowle et al. (2017) showed how SVs on neo-sex chromosomes drive hybrid male sterility, demonstrating that chromosomal rearrangements can underpin the genetic basis of speciation via genes directly involved in fertility. We also observed SVs impacting genes linked to olfactory function (Table S2), similar processes were identified in the SNP based approach of Kessler et al. (2024). Differences in olfactory genes might facilitate ecological partitioning between WTD and MD, contributing to their speciation. The role of olfactory genes in mate recognition and behavioral adaptation has been seen in house mice (North et al., 2020), where copy number variations in olfactory receptor genes influenced mate choice and reinforced reproductive isolation. This is particularly relevant in deer, as differences in olfactory responses might not only dictate ecological adaptations but also impact mate recognition, creating a dual mechanism for speciation through both environmental and sexual selection. Collectively, such fixed SVs and the corresponding phenotypic variation might implicate possible Dobzhansky-Muller incompatibilities in maintaining species boundaries in deer.

Further, several genes associated with species-specific SVs exhibited evidence of positive selection in both MD and WTD. The SV impacting genes with positive dN\dS values (3 total), were translocations and duplications, but less clear links to reproductive or ecological divergence (Table S2). Here we note one limitation in evaluating the evolutionary consequences of duplications lies in SV detection, as current SR-based tools identify but do not reconstruct the SV, with the absence of the SV in the assembly reflective of the common practice of purging duplications (Guan et al, 2020). As a result, dN/dS calculations are confined to the sequence of the initial (homologous) duplicated region, though LR approaches will allow for assess the evolution of duplicated regions.

### Differences between SV detection methods and future directions

The accurate estimation of SV size can be influenced by the tools and methods employed (Sedlazeck et al., 2018; Jeffares et al., 2017). The accuracy and sensitivity of SV callers vary significantly, as some tools are better suited for detecting larger SVs, while others excel at identifying smaller changes (Sedlazeck et al., 2018; Sudmant et al., 2015; Jeffares et al., 2017). This inherent bias across tools shapes the type of SV detected and the observed size distribution. To account for this, we employed two distinct data streams for SV detection to evaluate genome differences between MD and WTD, both reflective of commonly available datasets. Our comparison of SR and LR SV datasets showed differences that shape how structural variation is represented in genomic studies. As demonstrated in earlier benchmarking and evolutionary studies (Pei et al., 2024; Mahmoud et al., 2019), LR technologies detect substantially more SVs across nearly all classes, and our results reflect this same pattern (Tables 3 and 4). This discovery is expected, as long reads traverse repetitive and low-complexity regions where most SV breakpoints occur, allowing callers to resolve thousands of smaller and intermediate events that SR platforms fail to detect. In contrast, the SR datasets captured markedly fewer SVs across all classes and exhibited patterns consistent with well-known short-read constraints (Mérot et al, 2023). The large SR sample size and requirement of 100% SV detection in all samples (both LR and SR), might be overly stringent, but provided confidence in the observed calls, especially those overlapping between data sets (Table 5). While the absolute counts between SR and LR calls differed (Table 3 and 4), any non-overlapping SV can be presumed a false-positive or false-negative, mostly likely occurring in the SR data. The varied scenarios (Table 1) also allowed for assembly artefacts and singletons to be removed, where the only explanation for SVs detected only in either WTD2WTD or MD2MD comparison (scenarios 1 and 2) is that assembly differs from the conspecific resequencing data. Ultimately, the combined tally of SVs (Table 5), reflect high-confidence calls, and are likely evolutionarily relevant.

Future research can build on our findings through several approaches aimed at confirming the role of SVs in speciation. Mouse models would offer experimental approaches to validate, but analyzing known hybrids and backcrosses in deer should allow assessing the distribution and inheritance patterns of SVs. RNA-seq combined with chromatin accessibility assays could allow us to measure how SVs and the enhancer-rich regions alter gene regulation in both species. Lastly, expanding the scope to comparative analyses across other cervid species will further clarify whether similar classes of SVs repeatedly contributed to divergence and allow for reconstructing the ancestral state of species-specific SVs in *Odocoileus*.

## Supporting information

FaezehAzimiChetabi_Supplementary_Material_Revision2

## Acknowledgements

We thank Alan Cain, Steeve Côté, Anh Dao, Marco Festa-Bianchet, Brad Fulk, Eric Hoffman, Levi Jaster, Daniel Koelsch, Emily Latch, Brent Patterson, Joe Nocera, Don Stewart, Russell Easy, Jonathan Shaw, David Walter, and Jon Wheeler for providing samples. We thank two reviewers for providing valuable feedback.

## Data availability

All SR and some of LR sequencing samples are publicly available in the NCBI database under BioProject accession PRJNA830519. New LR data has been deposited (PRJA ID pending). And ortholog alignments in Figshare (https://doi.org/10.6084/m9.figshare.31830535). The code supporting this work is available in our lab’s GitLab repository: https://gitlab.com/WiDGeT_TrentU/graduate_theses/-/tree/master/AzimiChetabi/Chapter%20one

## Author Contribution

We had no external contributions.

