## Supplementary material for "Structural Genomic Variation And Its Potential Role In Deer Speciation": FaezehAzimiChetabi_Supplementary_Material_Revision2

**Figure S1.** A detailed plot of short-read workflow. Species are abbreviated as follows: MD = Mule deer, WTD = White tailed deer.

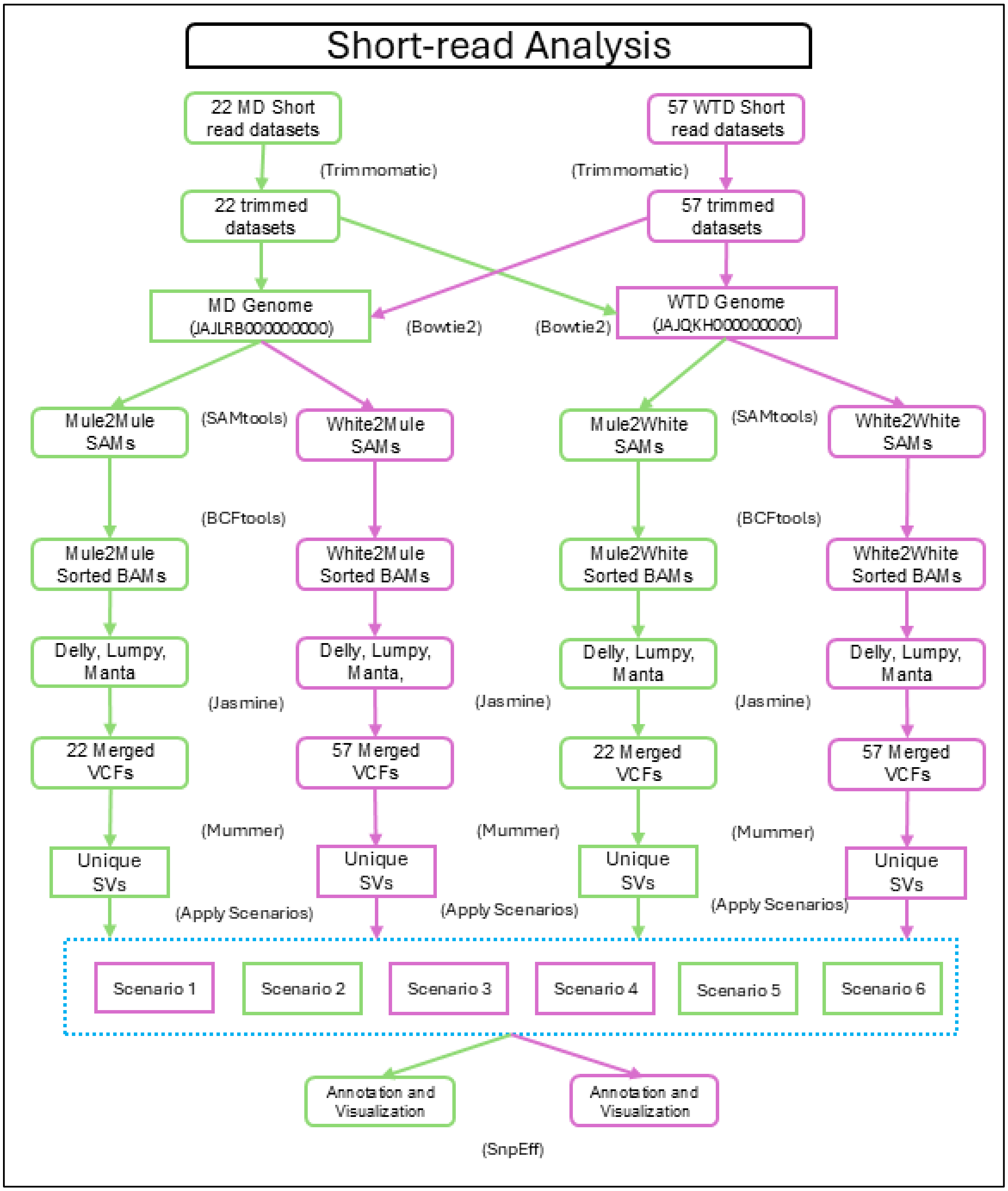

**Figure S2.** A detailed plot of long-read workflow. Species are abbreviated as follows: MD = Mule deer, WTD = White tailed deer.

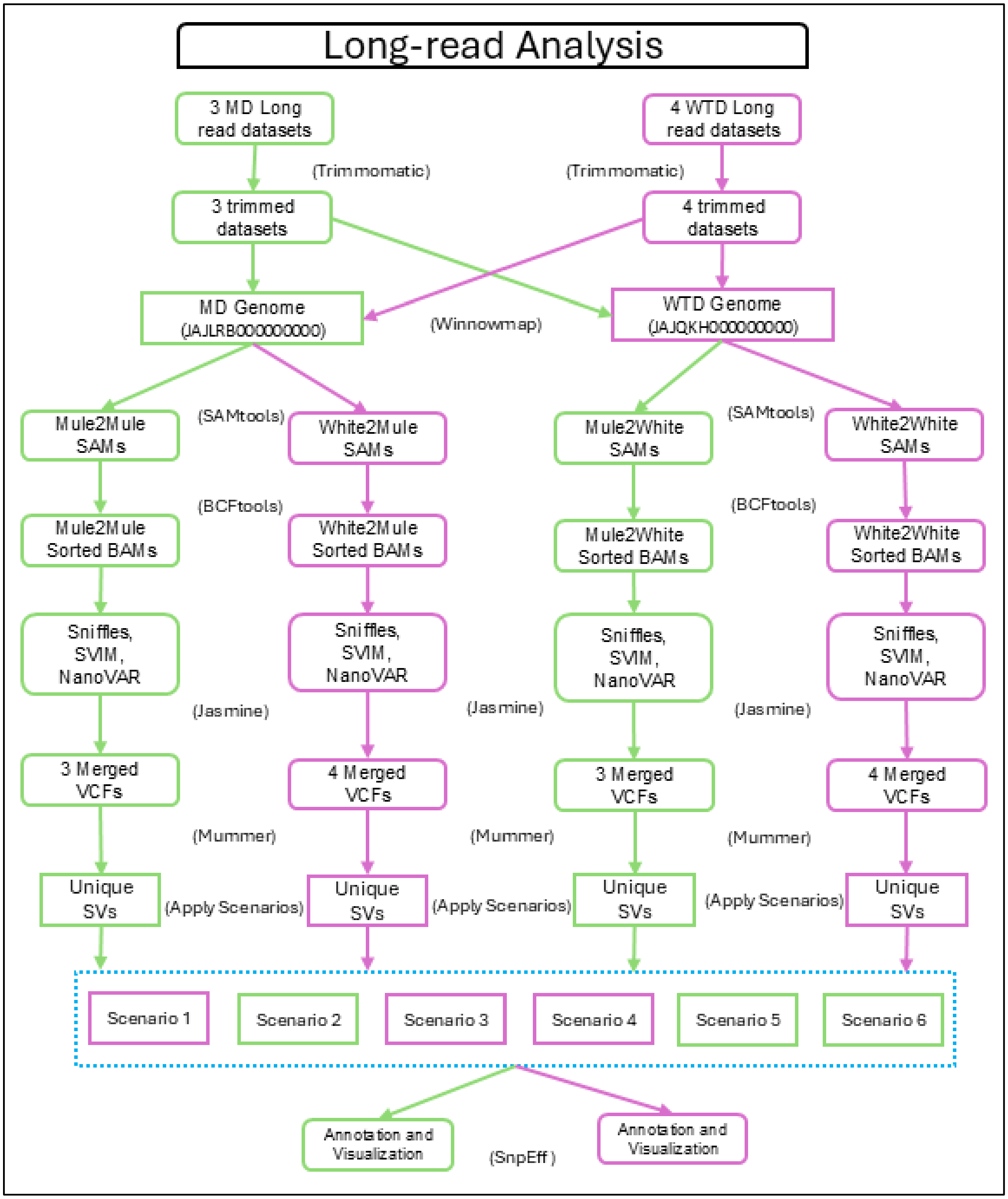

**Table S1.** Percentage of homozygous structural variants (SVs) detected in the short read and long read analyses (Figure S1 & S2) across all mapping strategies (MD2MD, MD2WTD, WTD2MD, WTD2WTD). For each SV type (deletion, duplication, insertion, inversion, translocation), the proportion of homozygous events was calculated as the number of homozygous SVs divided by the total SVs detected in that scenario.

| | MD2MD | MD2WTD | WTD2MD | WTD2WTD |
|---|---|---|---|---|
| | | **Short read SV analysis** | | |
| Deletion | 74.93 | 93.49 | 77.66 | 66.21 |
| Duplication | 6.17 | 3.28 | 1.42 | 1 |
| Insertion | 77.01 | 78.73 | 68.84 | 63.86 |
| Inversion | 24.94 | 44.22 | 24.65 | 20.92 |
| Translocation | 4.87 | 2.92 | 3.69 | 3.52 |
| | | **Long read SV analysis** | | |
| Deletion | 39.39 | 68.3 | 54.36 | 41.39 |
| Duplication | 28.65 | 27.07 | 35.4 | 29.18 |
| Insertion | 43.17 | 63.59 | 48.55 | 39.62 |
| Inversion | 25.93 | 58.17 | 37.5 | 37.56 |
| Translocation | 1.32 | 7.65 | 18.44 | 15.98 |

**Table S2.** Mule and white-tailed deer genes found within fixed species–specific structural variants and present in 100% of samples. Gene name from the white-tailed deer annotation with Orthogroup ID. Type of structural variant (SV). Significant P-values calculated by BUSTED (Murrell et all, 2015) and Bonferroni-Holm corrected are denoted by * on the Orthogroup ID. Citations for putative male reproductive and olfactory links of impacted genes are provided.

| Orthogroup ID | Gene Name | SV-type | Potential Speciation link |
|---|---|---|---|
| OG0016858 | MERTK | Translocation | Male reproduction - (Shi et al, 2023) |
| OG0016300* | CD70 | Translocation | - |
| OG0016712 | LOC110146842 | Translocation | Olfactory receptor – (NCBI, 110146842) |
| OG0017397 | KCNK1 | Translocation | Male reproduction and olfactory - (Delgado-Bermúdez et al, 2025; Yu et al, 2024) |
| OG0017682* | LRRC38 | Translocation | - |
| OG0017020 | OLFML2A | Translocation | - |
| OG0015447* | COL19A1 | Duplication | - |
| OG0016445 | FABP6 | Deletion | - |
| OG0017723 | TASL | Translocation | - |
| OG0014825 | IGFBP3 | Translocation | Male reproduction - (Fu et al, 2021) |
| OG0016518 | PAPPA2 | Translocation | Male reproduction - (Zamil et al, 2024) |
| OG0014866 | CREB3L2 | Duplication | - |
| OG0016712 | LOC110146842 | Inversion | Olfactory receptor – (NCBI, 110146842) |
| OG0015201 | TLCD1 | Translocation | - |
| OG0016445 | FABP6 | Translocation | - |
| OG0015643 | TAS1R2 | Inversion | Male reproduction and olfactory - (Mosinger et al, 2013; Zhao et al, 2010) |
| OG0014825 | IGFBP3 | Deletion | Male reproduction - (Fu et al, 2021) |
| OG0015470 | TUBE1 | Translocation | Male reproduction - (Stathatos et al, 2024) |
| OG0015643 | TAS1R2 | Translocation | Male reproduction and olfactory - (Mosinger et al, 2013; Zhao et al, 2010) |
| OG0014866 | CREB3L2 | Duplication | - |

| | | | |
|---|---|---|---|
| OG0015681 | SLC16A8 | Translocation | - |
| OG0017560 | EDNRB | Insertion | Olfactory - (Bryche et al, 2020) |
| OG0016858 | MERTK | Translocation | Male reproduction - (Shi et al, 2023) |
| OG0014916 | LOC110148892 | Deletion | - |
| OG0015201 | TLCD1 | Translocation | - |
| OG0016518 | PAPPA2 | Deletion | Male reproduction - (Zamil et al, 2024) |
| OG0013553 | RXRG | Deletion | Male reproduction - (Wang et al, 2020) |

**Table S3.** Motif density in mule deer (MD) and white-tailed deer (WTD) genomic regions (genic regions masked). Comparisons include gene bodies, ±50 kb gene flanking regions, 10,000 random SVs (±50 kb), and 27 fixed SVs (±50 kb). Final column reports the standard deviation of the average motif density values (Avg Motifs/bp).

| ***Data*** | ***Mean # Motifs*** | ***Avg Motifs / bp*** | ***Standard Deviation*** |
|---|---|---|---|
| *Genome (intergenic)* | 481.06 | 0.01099 | 0.01978 |
| *Genic regions +/- 50Kb* | 296.1475 | 0.00594 | 0.00540 |
| *Random SV +/- 50Kb* | 122.37 | 0.00246 | 0.00211 |
| *Fixed SVs +/- 50Kb* | 35.99 | 0.00078 | 0.00024 |
